# C->U transition biases in SARS-CoV-2 – still rampant four years from the start of the COVID-19 pandemic

**DOI:** 10.1101/2024.06.18.599635

**Authors:** Peter Simmonds

## Abstract

The evolution of SARS-CoV2 in the pandemic and post-pandemic periods has been characterised by rapid adaptive changes that confer immune escape and enhanced human-to-human transmissibility. Sequence change is additionally marked by an excess number of C->U transitions suggested as being due to host-mediated genome editing. To investigate how therse influence the evolutionary trajectory of SARS-CoV-2, 2000 high quality, coding complete genome sequences of SARS-CoV-2 variants collected pre-September, 2020, and from each subsequently appearing alpha, delta, BA.1, BA.2, BA.5, XBB, EG, HK and JN.1 lineages were downloaded from NCBI Virus in April, 2024. C->U transitions were the most common substitution during diversification of SARS-CoV-2 lineages over the 4-year observation period. A net loss of C bases and accumulation of U’s occurred at a constant rate of approximately 0.2%-0.25% / decade. C->U transitions occurred in over a quarter of all sites with a C (26.5%; range 20.0%-37.2%), around 5 times more than observed for the other transitions (5.3%-6.8%). In contrast to an approximately random distribution of other transitions across the genome, most C->U substitutions occurred at statistically preferred sites in each lineage. However, only the most C->U polymorphic sites showed evidence for a preferred 5’U context previously associated with APOBEC 3A editing. There was a similarly weak preference for unpaired bases suggesting much less stringent targeting of RNA than mediated by A3 deaminases in DNA editing. Future functional studies are required to determine editing preferences, impacts on replication fitness *in vivo* of SARS-CoV-2 and other RNA viruses and impact on host tropism.

## INTRODUCTION

The COVID-19 pandemic between 2020-2023 followed the emergence of severe acute respiratory syndrome coronavirus 2 (SARS-CoV-2) in 2019. SARS-CoV-2 is a member of the genus *Betacoronavirus* in the family *Coronaviridae* (1) and likely originates from a zoonotic spillover of a variant of sarbecoviruses widely distributed in *Rhinolophus* (horseshoe) bats in South-East Asia (2). Considerable insight into the evolution of SARS-CoV-2 over the pandemic period and beyond has been obtained through the large international sequencing effort, with over 16 million complete genome sequences catalogued to date. SARS-CoV-2 rapidly diversified into several genetically distinct clades (Fig. 1) and multiple cycles of emergence and extinction of variants of concerns (VoC), including alpha and delta, and their replacement by omicron in 2022. This has in turn diversified into several further lineages such as BA.1-5, XBB, and most recently JN.1.

**FIGURE 1.**
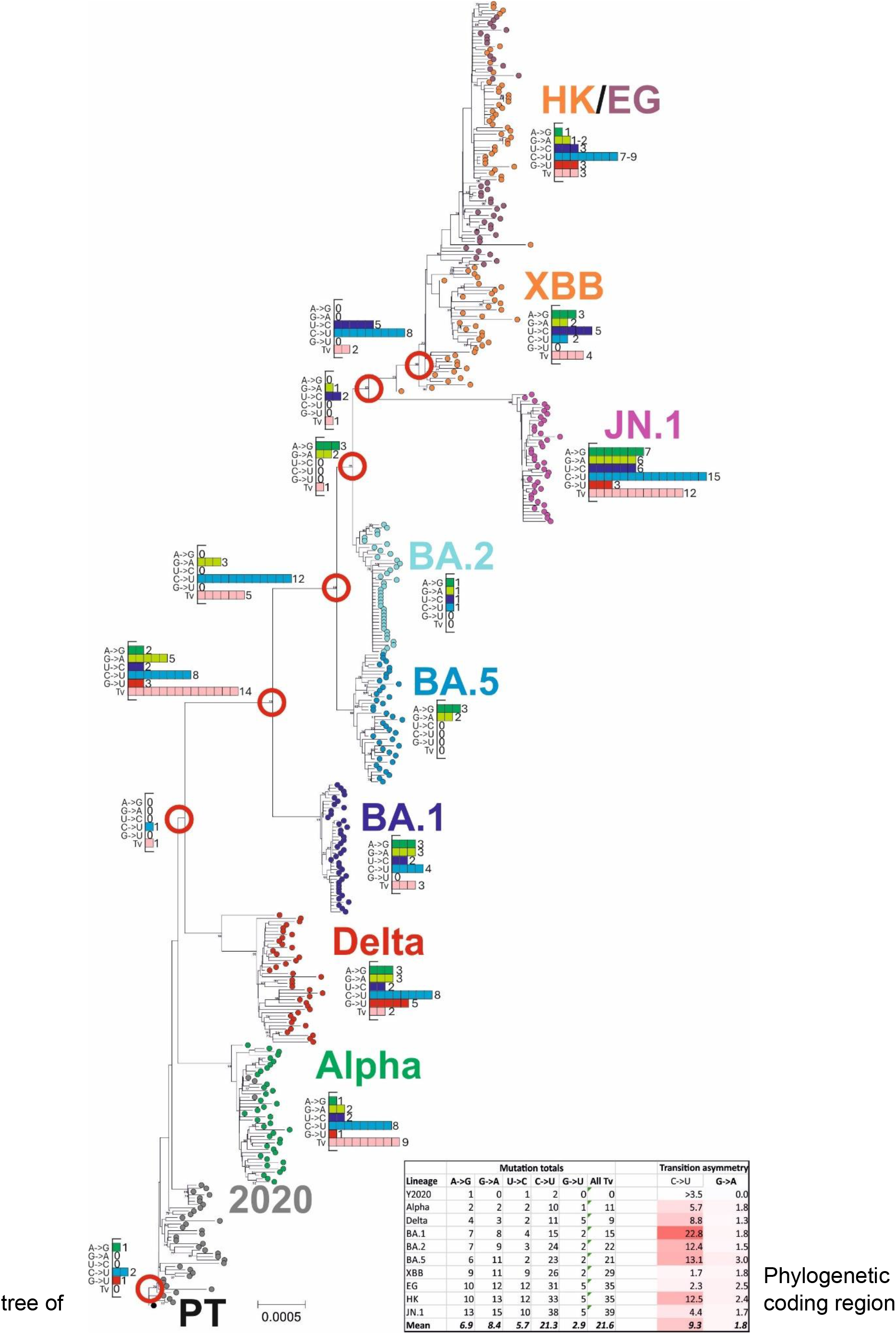
SEQUENCE CHANGES IN THE EMERGENCE OF SARS-CoV-2 LINEAGES. Phylogenetic tree of coding region sequences of lineages of SARS-CoV-2 sequentially emerging during the course of the COVID-19 pandemic during 2020-2024. Inset histograms show the numbers of each transition, G->U and total transversions that occurred between nodes (red circles) and descendant nodes or consensus lineage sequence. Substitution totals for each lineage are provided in the inset table, along with estimated transition asymmetries that account for base composition. The tree was constructed by neighbour joining of Jukes-Canrtor corrected distances; robustness of grouping was determined by boostrap re-sampling of 100 pseudo-replicas; support values of >70% are shown on the branches.

Sequence change in SARS-CoV-2 over this period has been characterised by the accumulation of neutral substitutions and rapid, phenotypically driven evolution of the spike gene. This has conferred major changes in antigenicity, neutralisation susceptibility and receptor binding, the latter potentially contributing to the marked increases in SARS-CoV-2 transmissibility over time, such as the D614G mutation (3, 4). It also became very rapidly apparent that SARS-CoV-2 isolate consensus sequences contained a large number of often transient C->U substitutions distributed throughout the genome (5–11). Although without direct functional evidence at the time, these were frequently speculated to originate from host-mediated RNA editing pathways that constitute part of the cellular innate immune response to virus infections. Of these, C->U substitutions are characteristic of the activity of members of the apolipoprotein B mRNA-editing enzyme, catalytic polypeptide-like (APOBEC) family (12). This is distinct from the activity of adenosine deaminase acting on RNA type 1 (ADAR1)(13) that targets double-stranded (ds) RNA replication complexes of RNA viruses. ADAR1 introduces A->G mutations into the virus genome as well as U->C mutations through editing of the complementary RNA strand in its dsRNA template.

The extent to which host-induced mutations contribute to the evolutionary trajectory of SARS-CoV-2 can be better investigated several years after the start of the pandemic. In the current study, the overlay of potentially host-directed editing of SARS-Cov-2 genomes with their diversification into distinct lineages has been investigated using a large dataset of sequences representing the principal SARS-CoV-2 lineages (alpha, delta, BA.1, BA.2, BA.5, XBB, EG, HK and JN.1) spanning the pandemic period to the present (April, 2024). As lineages each have a likely clonal origin, analysis of unfixed diversity within each lineage enables independent observations of site heterogeneity, providing a powerful method to investigate the potential existence of favoured targets for mutation and the contexts in which they occur.

## MATERIALS AND METHODS

### Sequence datasets

New datasets of SARS-CoV-2 were constructed from publicly available complete genome sequences on GenBank, selected using NCBI Virus for lineage, genome sequence quality and completeness (www.ncbi.nlm.nih.gov/labs/virus/). All available SARS-CoV-2 variants assigned as Variants of Concern (VoCs) alpha, and delta and the omicron lineages BA.1, BA.2, BA.5, XBB, EG, HK and JN.1 were downloaded on the 12^th^ April, 2024. A further dataset of early SARS-CoV-2 variants not assigned to a lineage was created from sequences with sample dates pre-September, 2020 (Y2020). From these downloads, 2000 sequences from each were randomly selected (only 1685 and 446 sequences of EG and HK were available) for analysis. GenBank annotations were used to assign sample dates to each sequence. Sequences were aligned by Nextclade (14), including the Wuhan-Hu-1 (MN908947/CN/2019) prototype strain in each alignment as a reference sequence.

A published compilation of substitutions observed within a collection of around 7 million SARS-CoV-2 genome sequences (including the set of all sequences in GISAID as of 29 March 2023; (15)) was used as an alternative data source. Sites were categorised by their relative frequency of occurrence, calculated as the natural logarithm of the ratio of actual to expected counts (described as the estimated fitness, δ*f*).

For both datasets, the regions spanning the start of the first open reading frame (ORF1a) to the end of the last ORF (NS10) were used for standard analysis to avoid areas of reduced coverage and potentially greater read error in the genome terminal regions.

Available full genome sequences from SARS-CoV-2 variants infecting deer (n=200) and mink (n=199) were generated similarly.

### Sequence analysis

Calculation of nucleotide composition was performed using the SSE package version 1.4 (16). Sequence changes of each full dataset (up to 2000 genome sequences) were compiled using the program SequenceChange; unfixed mutations were identified through the use of a <5% site variability threshold, calculated as the cumulative frequency of all non-consensus bases. Raw data on substitution frequences at each genome position used in the analysis are provided in Table S1; Suppl. Data.

As described previously (5), normalised transition asymmetries of C->U and U->C substitutions (and comparably for G->A and A->G) were calculated as f(C->U) / f(U->C) * (fU/fC), where f = frequency.

Potential biases in the identities of bases immediately 5’ and 3’ to C->U mutated sites was performed by a new modelling approach. This was based on calculation of observed frequencies of bases either side of C->U mutated sites in the concatenated ORF1a/ORF1b reading frame within alignments of representative sequence lineages (Y2020, Delta, BA.1 and JN.1; 1000 sequences in each) spanning the observation period of the study. These frequencies were then compared with those of an equivalent sized dataset of sequences that had been randomised in sequence. The program SequenceMutate in the SSE package performed sequence randomisation under constraints – NDR: dinucleotide sequences preserved, COR: coding preserved; CDLR: coding sequence and dinucleotide frequencies preserved.

### Phylogenetic analysis

Neighbour joining trees were constructed from aligned sequences using the program MEGA7 (17).

### Statistical analysis

All statistical calculations and histogram constructions used SPSS version 29.

## RESULTS

### Sequence change during lineage evolution

Datasets of 2000 aligned sequences of SARS-CoV-2 were selected to represent early pandemic onset strains (Y2020; pre-September, 2020), VoCs alpha and delta, and the omicron lineages BA.1, BA.2, BA.5, XBB, EG, HK and JN.1. These emerged sequentially in the four years since the start of the COVID-19 pandemic. Base substitutions associated with the emergence of each lineage were determined though enumeration of sequence differences of lineage consensus sequences with the prototype Wuhan-1 strain (Fig. 1).

Substitutions from the prototype SARS-CoV-2 sequence were characterised by an excess of C->U substitutions (light green) over other transitions and transversions throughout the tree.

Major branching points were additionally associated with higher proportion of transversions (pink bars) resulting from selected amino acid changes in the evolution of the spike gene. This was confirmed by analysis of the positions of each substitution type (Fig. 2). While there was a marked predominance of C->U changes in the ORF1a/1b gene and between ORF3a and the end of ORF10, there was a much higher proportion of transversions in the spike gene with well characterised involvement in antigenic change.

**FIGURE 2.**
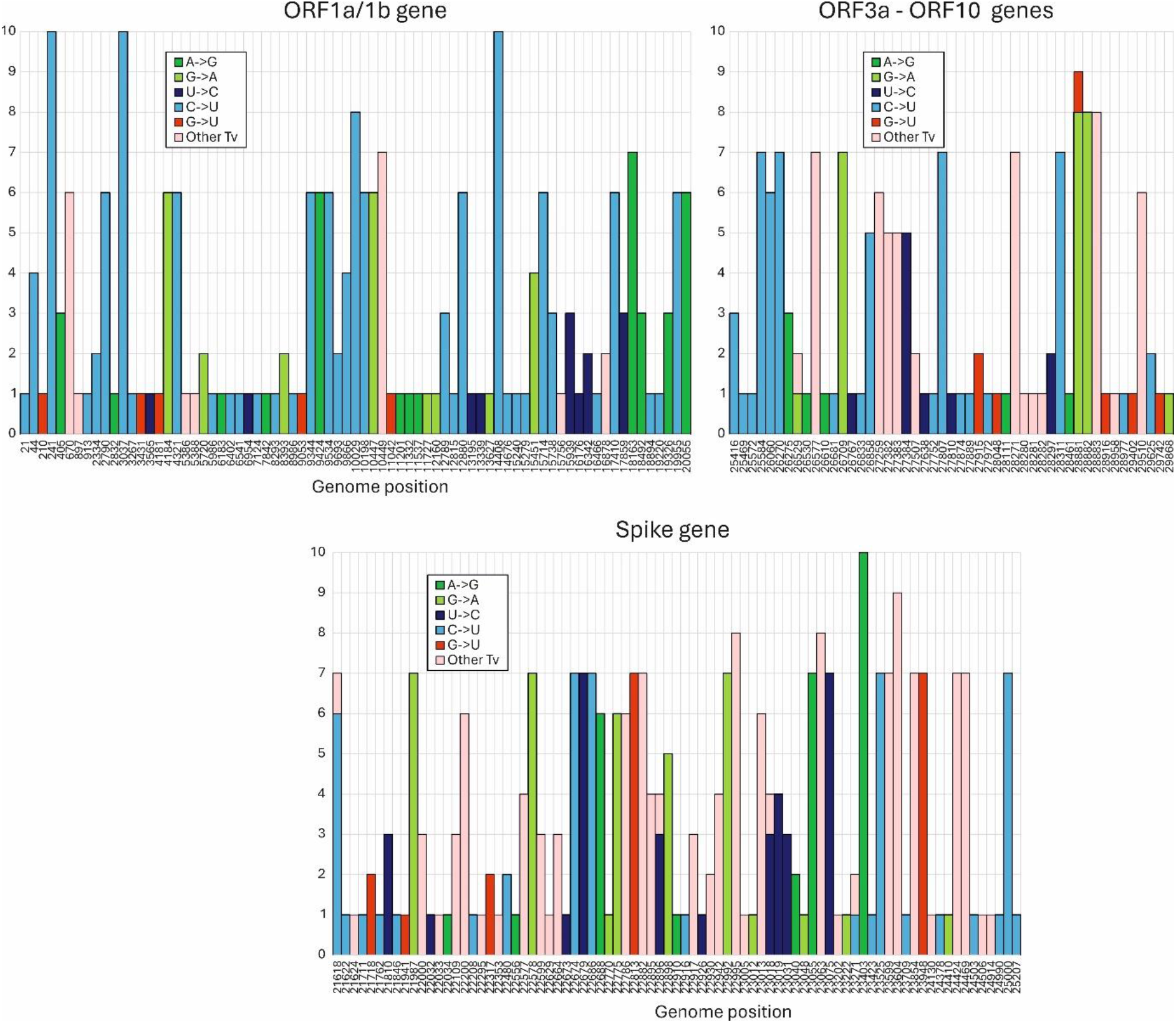
GENOME POSITION OF SUBSTITUTIONS IN SARS-CoV-2 LINEAGE CONSENSUS SEQUENCES. Positions of mutations in consensus sequences in the SARS-CoV-2 genome (only variable sites are plotted; positions indicated on the x-axis). The y-axis records numbers of lineages containing each indicated mutation. For clarity, the genome has been divided into ORF-1a/1b, S gene and ORF3a-ORF10 sub-regions.

The effects of mutational biases in SARS-CoV-2 genomes on their overall composition was investigated by plotting C and U mononucleotide frequencies of the 10 lineage consensus sequences with mean sampling dates calculated from their component SARS-CoV-2 genome sequences (Fig. 3A, 3B). There was a progressive decline in the frequency of C and a corresponding rise in the frequencies of U consequent to the accumulated C->U transitions that was strongly coupled to sampling date (R^2^ vales of 0.66 and 0.64 respectively). Linear trendlines converged on the base composition of Wuhan-1 around the end of 1989. C and U had trajectories in depletion and accumulation of 0.2% -0.25% / decade (Fig. 3C). Much smaller changes in the composition of G and A were observed (0.05% – 0.06%), reflecting the much lower G->A/A->A transition asymmetry compared to C->U/U->C (Fig. 1); the depletion of G may additionally reflect previously observed higher rate of G->U transversions (8, 10).

**FIGURE 3.**
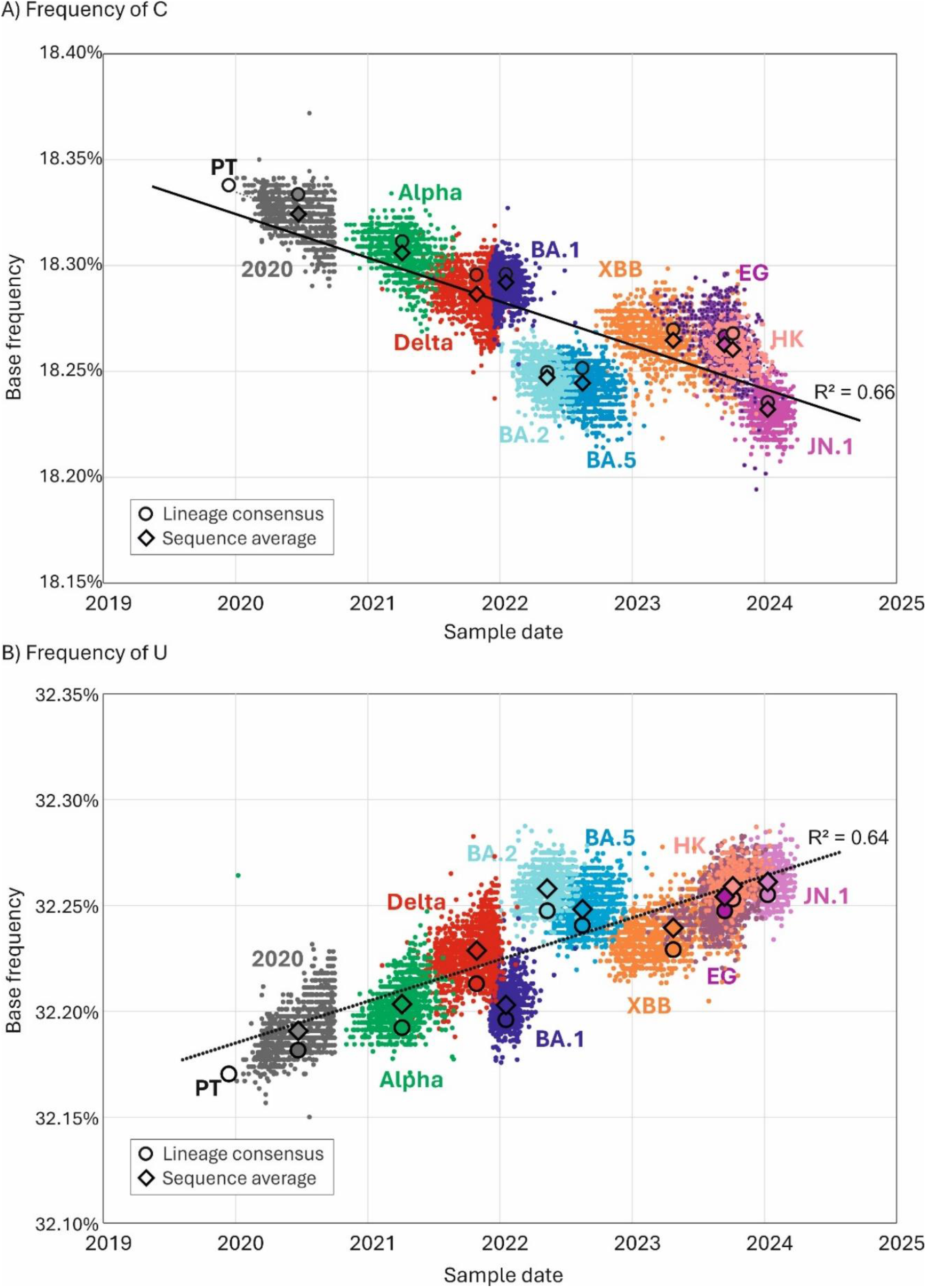
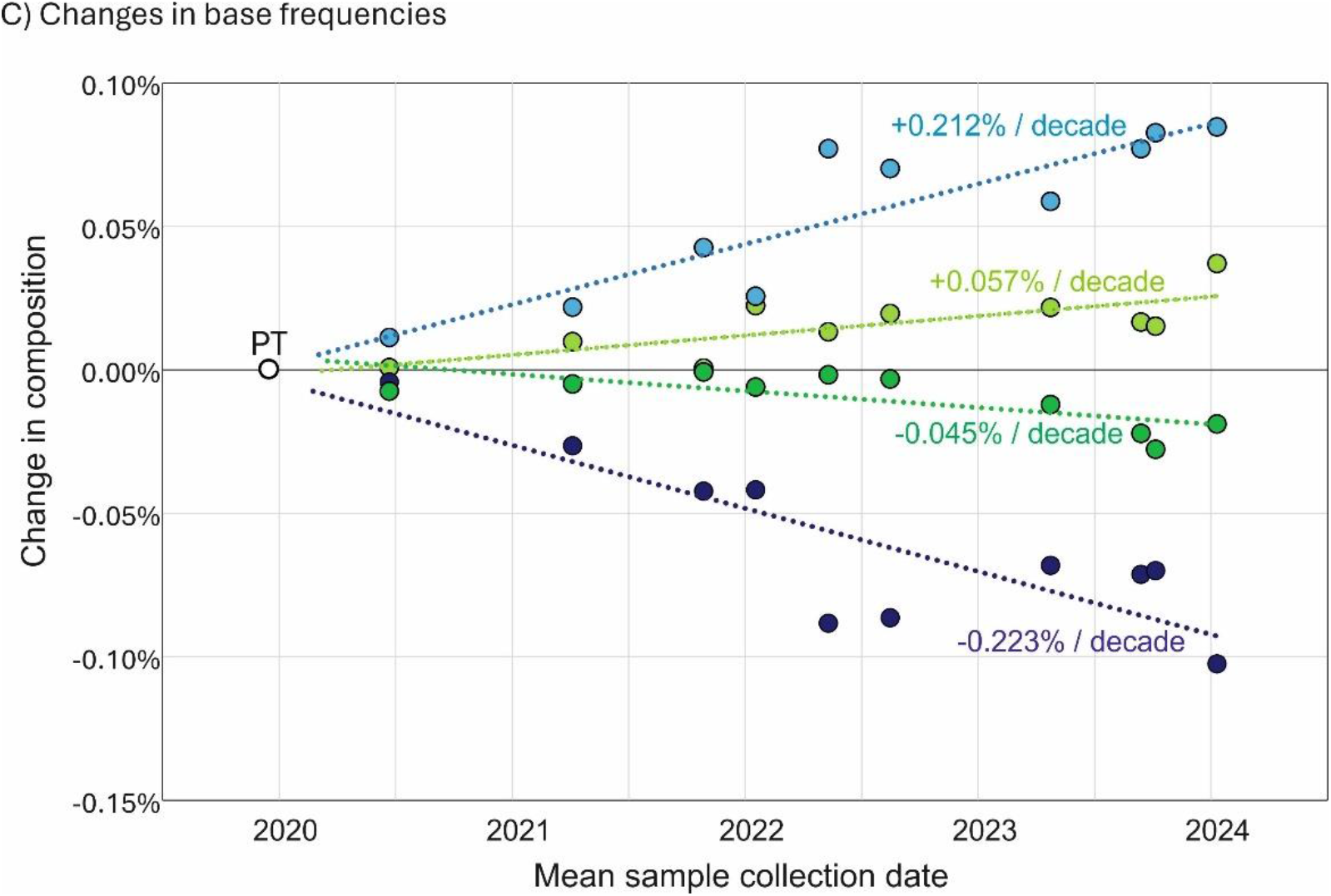
CHANGES IN SARS-CoV-2 GENOME COMPOSITION SINCE THE START OF THE PANDEMIC. (A, B) Frequencies of C and U bases in individual genome sequences of SARS-CoV-2 isolates collected at different time-points from the start of the pandemic. Lineage assignments are indicated. Circles indicate mean sampling date and composition of consensus sequences from the 10 lineages and the prototype sequence; diamonds inidicate mean compositions of individual sequences. (C) differences in A, C, G and U base frequencies between the genome sequence of the PT (Wuhan-1) and of lineage consensus sequences collected at different time points from the start of the pandemic.

Compositional biases in other coronaviruses were consistent with longer term effects of the C->U transition asymmetry (Fig. 4). All four human seasonal coronaviruses in the *Alphacoronavirus* (HCoV-229E and HCoV-NL63) and *Betacoronavirus* (HCoV-OC43 and HCoV-HKU1) genera showed far greater imbalances of C/G and U/A base frequencies (G/C: 4.9%-6.1%; U/A: 7.6%-12.8%) than found in SARS-CoV-2, SARS-CoV or any of the sarbecoviruses infecting bats (G/C: 0.93%-2.2%; U/A: 2.1%-3.4%). The extent to which this reflects a greater accumulated C->U mutational effect in non-bat hosts is discussed below.

**FIGURE 4.**
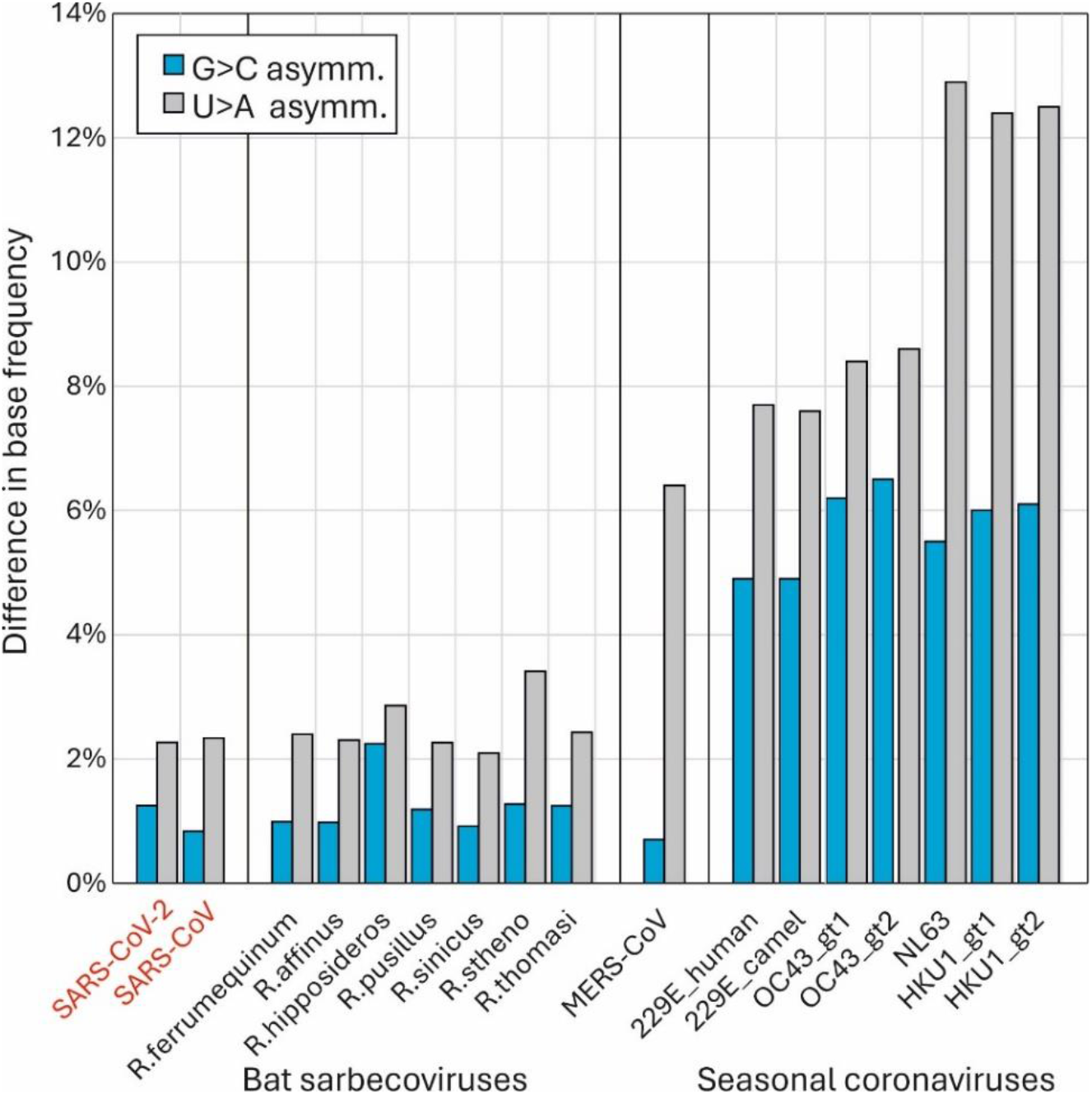
COMPOSITIONAL ASYMMETRIES IN OTHER CORONAVIRUSES Comparison of base compositions of the recently zoonotic SARS-CoV-2 and SARS-CoV isolate sequences with sarbecoviruses infecting the *Rhinolophus* genus of bats (host species indicated on x-axis) and with genotypes of proposed potentially zoonotically acquired human seasonal coronaviruses. Blue bars record the excess of G over U (U depletion), and grey bars the excess of U over A (U accumulation).

### Within-lineage base substitutions

Since the lineages analysed in the study have a primarily clonal origin, sequence diversity within lineages arises from evolutionarily independent events occurring within each. Furthermore, enumerating sites where mutation frequencies are less than 5% enables an inference of directionality (as previously discussed; (18)) that is independent of the evolutionary reconstructions used in the analysis of consensus sequences in the previous section. The use of large datasets of sequences for each lineage (typically 2000 whole genome sequences) revealed around a thousand times more polymorphic sites than were apparent from parsimony-based sequence reconstructions. As they were unfixed, this also provides the means to investigate the characteristics of C->U and other mutated sites using ten evolutionarily independent sets of observations from each lineage.

Applying the 5% heterogeneity filter to record unfixed substitutions within each lineage and their directionality, each showed similar excesses of C->U transitions compared to other substitutions (Fig. 5). The range of values (4.1-9.9) encompassed the mean transition asymmetry observed in consensus sequences (9.3). Contrastingly, there was a virtual absence of G->A/A->G transition asymmetry (ratios of 1.0-1.49) in any of the lineages.

**FIGURE 5.**
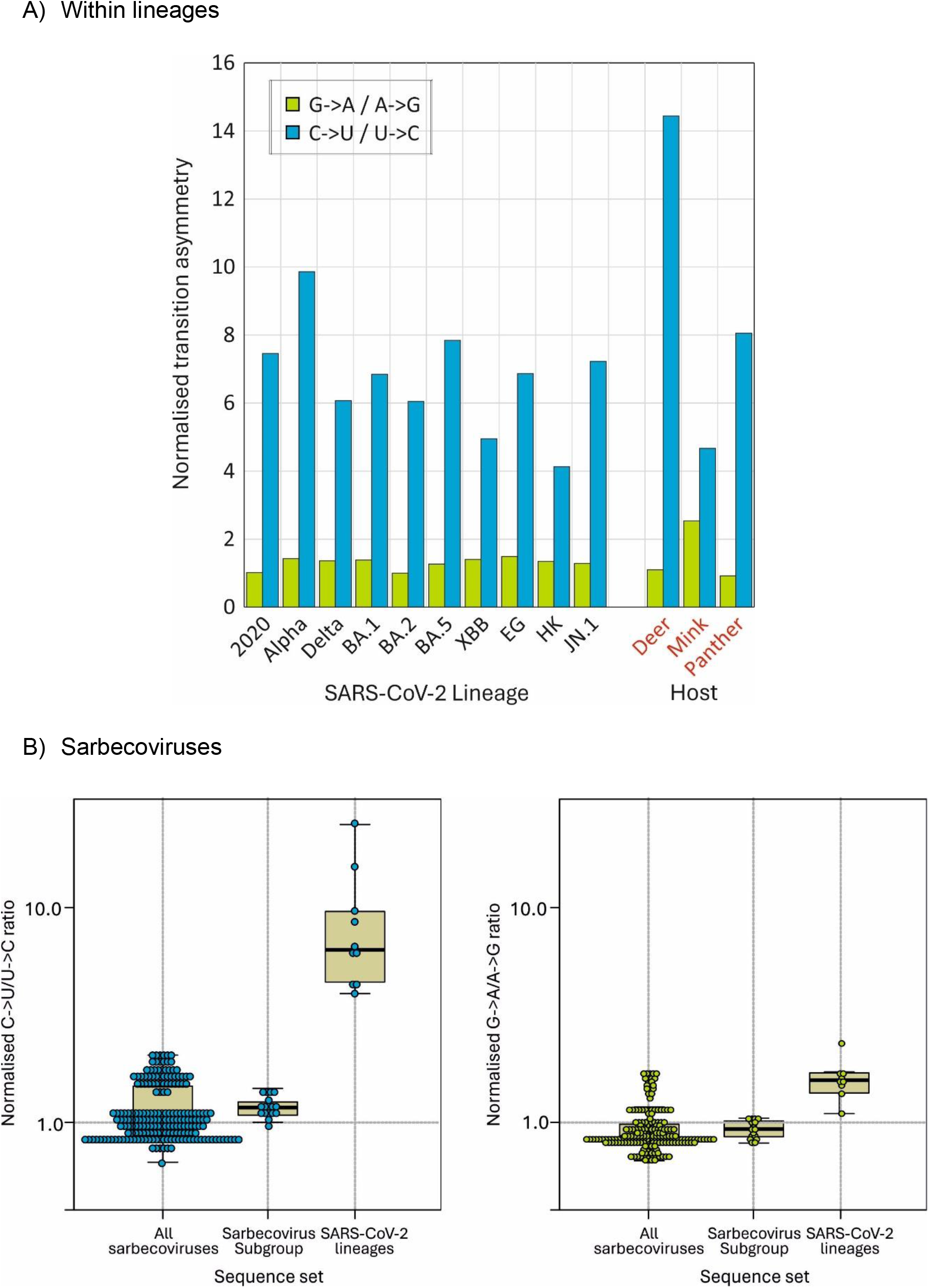
TRANSITION ASYMMETRIES WITH SARS-CoV-2 LINEAGES. A) Normalised transition asymmetries of fixed C->U and G->A substiutions within the 10 SARS-CoV-2 lineages. Comparisons with SARS-CoV-2 strains infection non-human species shown on righ panel. (B) Normalised transition asymmetries of all sarbecovirus strains infecting bats (column 1), the subset (including RatG13) most closely related to human SARS-CoV-2 (column 2) and consensus seuqences of lineages within SARS-CoV-2 compared to the prototype Wuhan-1 isolate sequence (column 3).

Transition asymmetries in SARS-CoV-2 variants infecting mink and panther were similar to those observed in human-derived variants, but much greater in deer (C->U/U->C: 14.4; G->A/A->G: 1.1). Transition asymmetries were characteristic of human and derived strains in animals, but not in sarbecoviruses more generally (Fig. 5B). There was no evidence for C->U/U->C (or G->A/A->G) asymmetries between the whole dataset of bat sarbecoviruses with a reconstructed ancestral sequence and with a subset of more closely related sequences variants including RatG13 and SARS-CoV-2.

Overall, discounting the datasets with fewer than 2000 available sequences (HK, EG), lineages showed a mean of 1429 sites (range 1075-2004) with C->U transitions, compared to sites with other transitions (A->G: 465; G->A: 392 and U->C: 534). Sites of C->U transitions occurred in over a quarter of all sites with a C (26.5%; range 20.0%-37.2%), around 5 times more than observed at sites with the appropriate majority base for the other transitions (5.3%-6.8%). To investigate whether certain sites were more likely to mutate than others, the distribution of sites with zero, 1, 2 or more lineages with C->U and other transitions was plotted and compared with the null expectation of a random distribution (Fig. 6). A-G, G->A and U->C transitions showed frequencies among lineages that closely matched an unbiased distribution calculated from the Poisson distribution based on mean frequencies of transitions per nucleotide position. However, the distribution of C->U transitions differed strikingly, with far more transition sites shared among 6 or more lineages, and conversely, an over-representation of invariant sites compared to the Poisson nul expectation. Sites of C->U substitutions in 3 or more lineages were consistently in large numerical excess to those with other transitions (total and proportions shown above graph). Favoured sites for C->U editing by A3A during *in vitro* passage of SARS-CoV-2 were recently reported (19). Most but not all of the 10 mutated sites were also those with high frequencies of unfixed C->U mutations among native SARS-CoV-2 variants (Fig. 6; Fig. S1, Suppl. Data), indicating some commonality in targeting (see Discussion).

**FIGURE 6.**
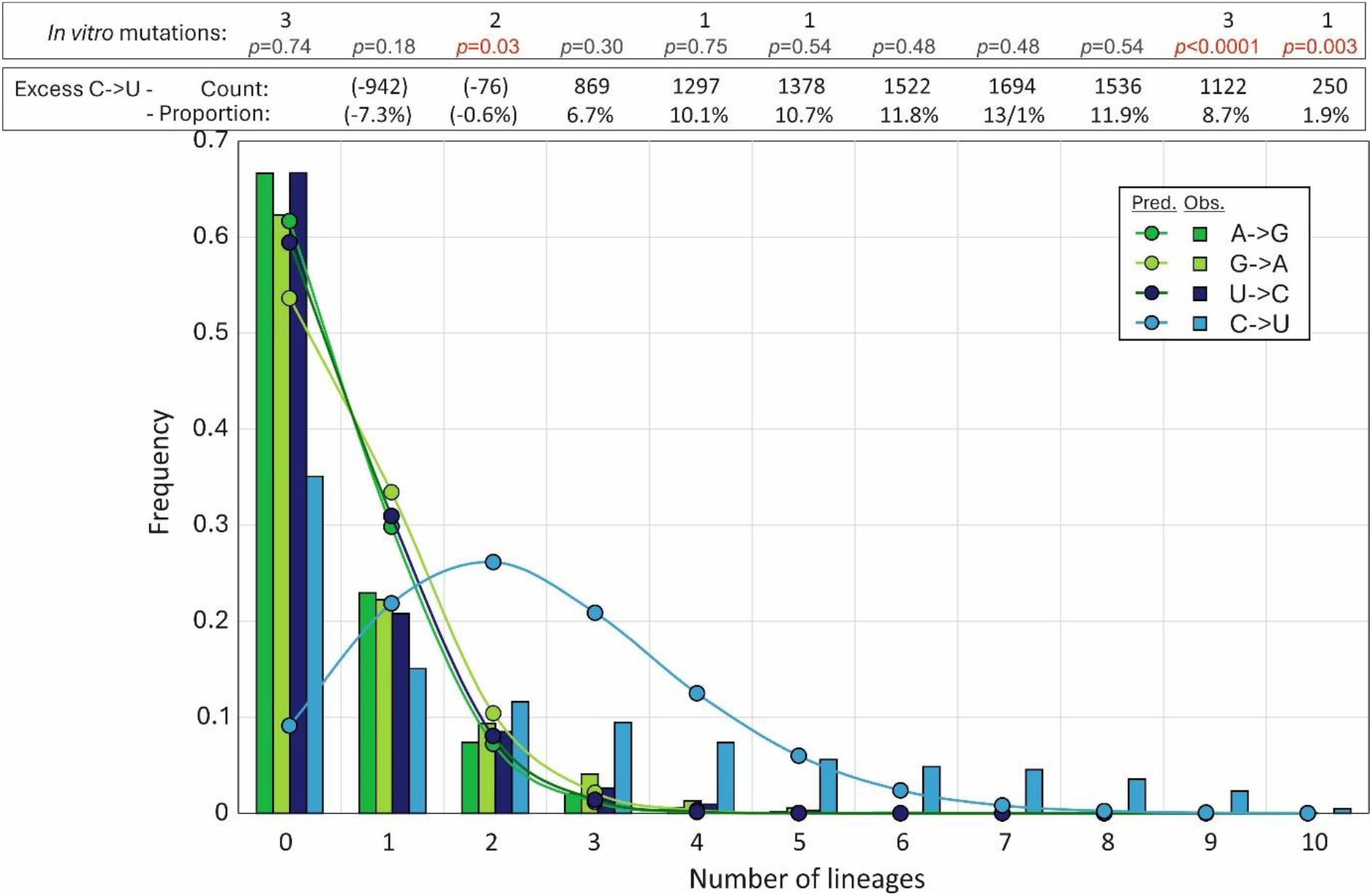
DISTRIBUTION OF MUTATIONS. The distribution of unfixed transitions (<5% frequency) between lineages (histogram) compared to expected values assuming no positional bias derived from the Pioisson distrribution (graph points). Excess numbers over the mean numbers of A->G, G->A, and U->C transitions (total excess 8651 from 12,907 C->U transitions) and their proportions of total C->U substitutions in each category are shown in the upper box. The distribution of C-U transitions at sites of previously reported *in vitro* A3A-induced mutations (19) is shown in the lower box (Fig. S1; Suppl. Data); differences in lineage distributions with observed values calculated by Pearson chi-square.

### Favoured contexts for C->U transitions

To investigate whether particular 5’ or 3’ bases were found in association with C->U substitutions, as typically found in sites edited by APOBECs, a novel modelling approach was developed to calculate expected frequencies based on a null expectation of no context bias. A more sophisticated approach was required beyond simply predicting frequencies from mononucleotide composition, as almost all sites lie within functional protein-encoding genes that imposes coding constraints from amino acid usage. Biases may additionally arise from biological selection against certain dinucleotides, such as CpG and UpA that may additionally impact upstream and downstream expected base composition.

To calculate this, expected frequencies of A, C, G and U upstream and downstream of a C residue were calculated from in-frame alignments of the ORF1a/ORF1b concatenated gene of four representative lineages (Y202, Delta, BA.1 and JN.1 spanning the observation period) using simple mononucleotide frequencies (29.9%, 18.3%, 19.6% and 32.2% respectively; Fig. 7A, 7B). However, observed frequencies of the four bases in native sequences of the four lineages differed markedly, irrespective of whether the C site was non-mutated, mutated in at least one lineage or in multiple lineages (red bar; Fig. 7A, 7B).

**FIGURE 7.**
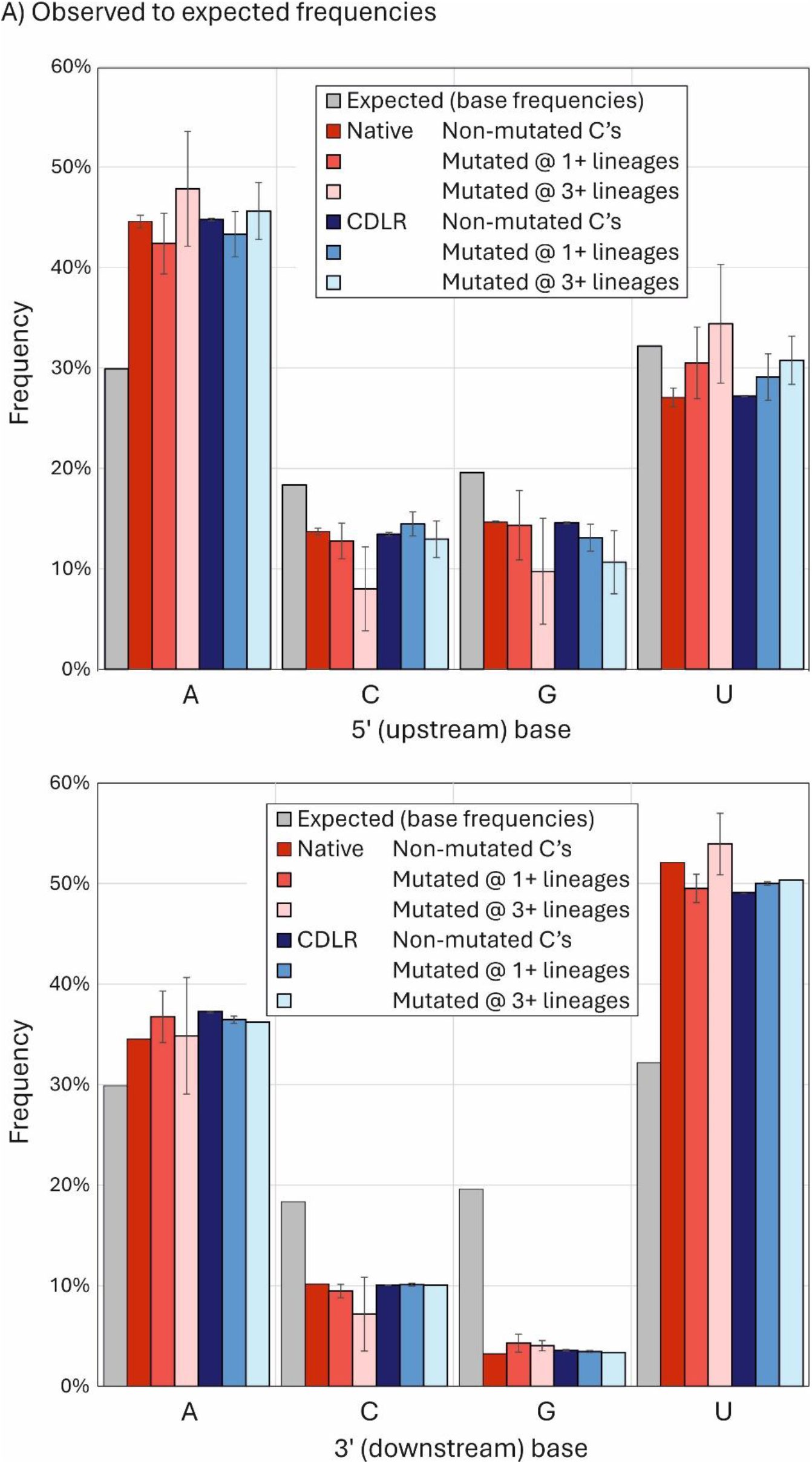
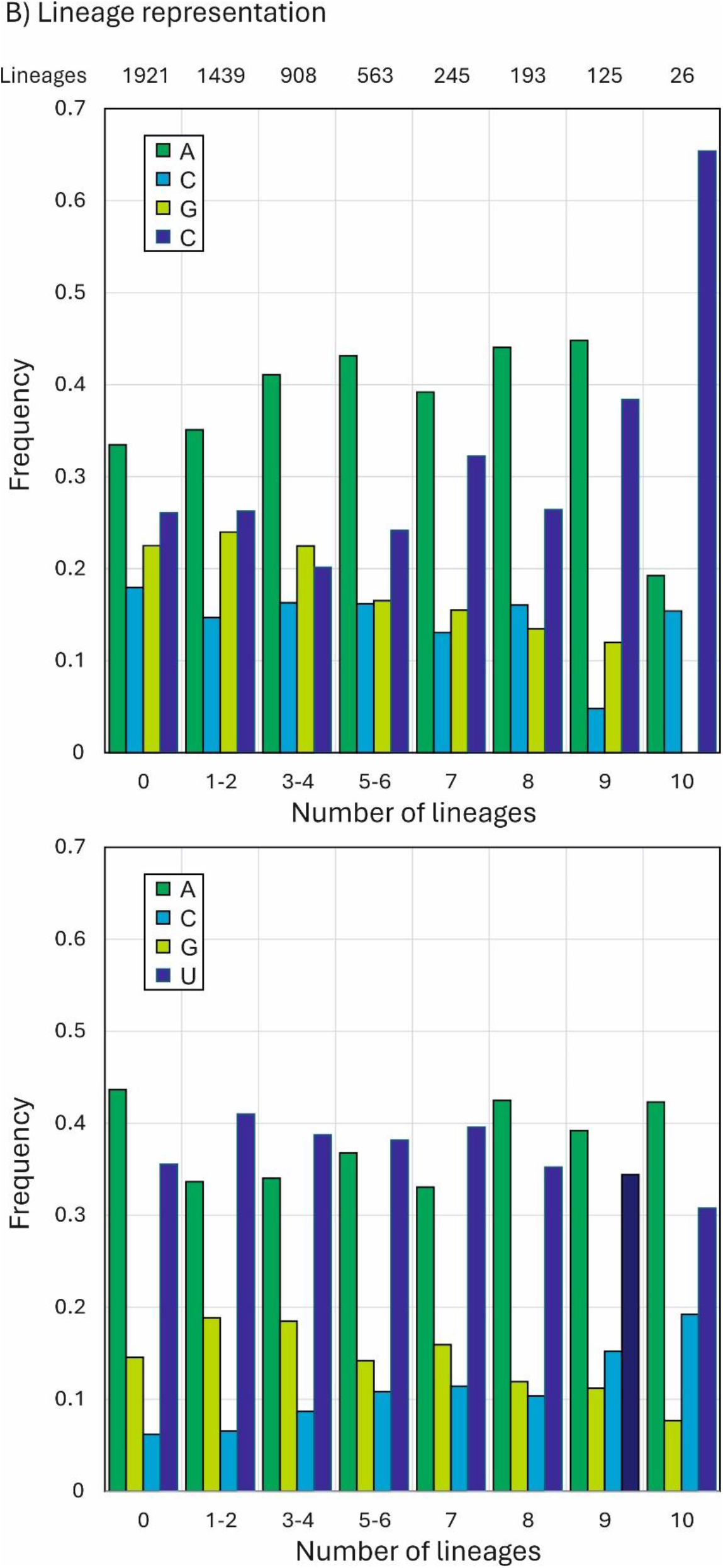
OBSERVED AND EXPECTED 5’ AND 3’ BASE CONTEXT FREQUENCIES AT C->U MUTATED SITES. (A) Predicted and onserved frequencies of bases5’ (upstream) and 3’ (dowenstream) of C’s, split into sites that were invariant or mutated in one or more lineages. Observed frequencies were compared with those of sequences randomised by CDLR that preserved native coding and dinucleotide frequencies. Frequency estimations based on mononucleotide frequencies alone are indicated by grey bars. Error bars for native and CDLR sequences show one standard deviation of values from the four lineages analysed (Y2020, delta, BA.1 and JN.1). (B) Distribution of 5’ and 3’ bases at C->U mutation sites occurring in different numbers of lineages and at invariant C sites.

Observed frequencies of upstream A of 44.6%, 42.4% and 47.8%) were significantly higher than the predicted 29.9%; similarly, frequencies of downstream U were substantially higher (52.1%, 49.5% and 53.9%) than the predicted 32.2%.

To determine whether these differences arose from favoured contexts for C->U mutations or whether these were expected frequencies once coding constraints and dinucleotide frequency biases in native sequences had been taken into account, a total of 1000 ORF1a/ORF1b sequence randomisations were performed on the consensus sequences of each of the four lineages. Randomisation using the algorithm CDLR (in the SSE package) created highly divergent sequences from the native sequence while preserving the protein coding of each as well as native dinucleotide frequencies. As encoded amino acid sequences were invariant, 3’ composition comparisons were most usefully performed at the downstream (+1) site of all C’s at codon position (CP)2 (all codons with a C at position 2 are 4-way redundant at CP3). Similarly, upstream (-1) base frequencies with and without sequence randomisation were compared for C’s at CP1 at CP3 of the preceding codon. For comparability, only 4-way redundant upstream codons were included in this comparison.

The analysis demonstrated virtual equivalence in both upstream and downstream base context frequencies between native SARS-CoV-2 sequences and those randomised by CDLR (Fig. 7A). As expected, the observed frequency of G downstream of C was much lower than predicted based on mononucleotide frequencies, reflecting the suppression of CpG in vertebrate RNA viruses; however, this bias was correctly reproduced in sequences scrambled by CDLR. Furthermore, there was no 5’ or 3’ compositional difference between C’s that were invariant from those that showed C->U changes in one or more lineages. All frequencies thus matched closely the null expectation once protein coding and dinucleotide biases had been taken into account.

While upstream and downstream base frequencies at C->U mutated sites were unbiased, most sites analysed in Fig. 7A were variable in 3 or fewer lineages. To investigate whether more frequently mutated C bases showed a more obvious 5’ or 3’ base context bias, sites were binned into 7 variability categories along with invariant sites, and 5’ and 3’ base frequencies recalculated (Fig. 7B). Remarkably, C->U mutation sites that occurred in 9 and 10 lineages showed an increasing and ultimately extreme 5’ context preference for U, while the 3’ context remained unchanged from less variable sites. However, there was a substantial over-representation in sites that polymorphic in between 5 and 8 lineages, but these failed to show a site preference for a 5’U (Table S2A; Suppl. Data). This indicates that the majority of sites over-represented for C->U substitutions sites did not possess the correct upstream base associated with A3A targeting.

### Association of C->U mutations with RNA secondary structure formation

A consensus RNA secondary structure model of the SARS-CoV-2 genome based on three independent studies using biochemical mapping and prediction methods (20–22) was used to determine whether C->U or other transitions occurred preferentially at sites that were non-base paired. The number of lineages with mutations occurring at unpaired or paired sites was compared with unpaired / paired ratios of invariant A, C, G and U bases (Fig. 8A). Ratios around 1 indicated no mutational bias towards paired or unpaired sites, and characterise the preferences of A->G, G->A and U->C transitions. Contrastingly, U->C and G->U mutations were strongly conditioned by pairing status, with a remarkable relationship between occurrence in multiple lineages and increasing mutational bias towards unpaired bases. For example, the 175 sites of C->U mutations occurring within 8 or more of the 10 SARS-CoV-2 lineages, 154 were unpaired compared to 25 paired (ratio 6.2), quite different from the unpaired/ paired ratio of invariant C sites (688 and 1311; ratio 0.53). There was consequently a >11.7-fold skew towards unpaired bases at highly mutated C-> sites.

**FIGURE 8.**
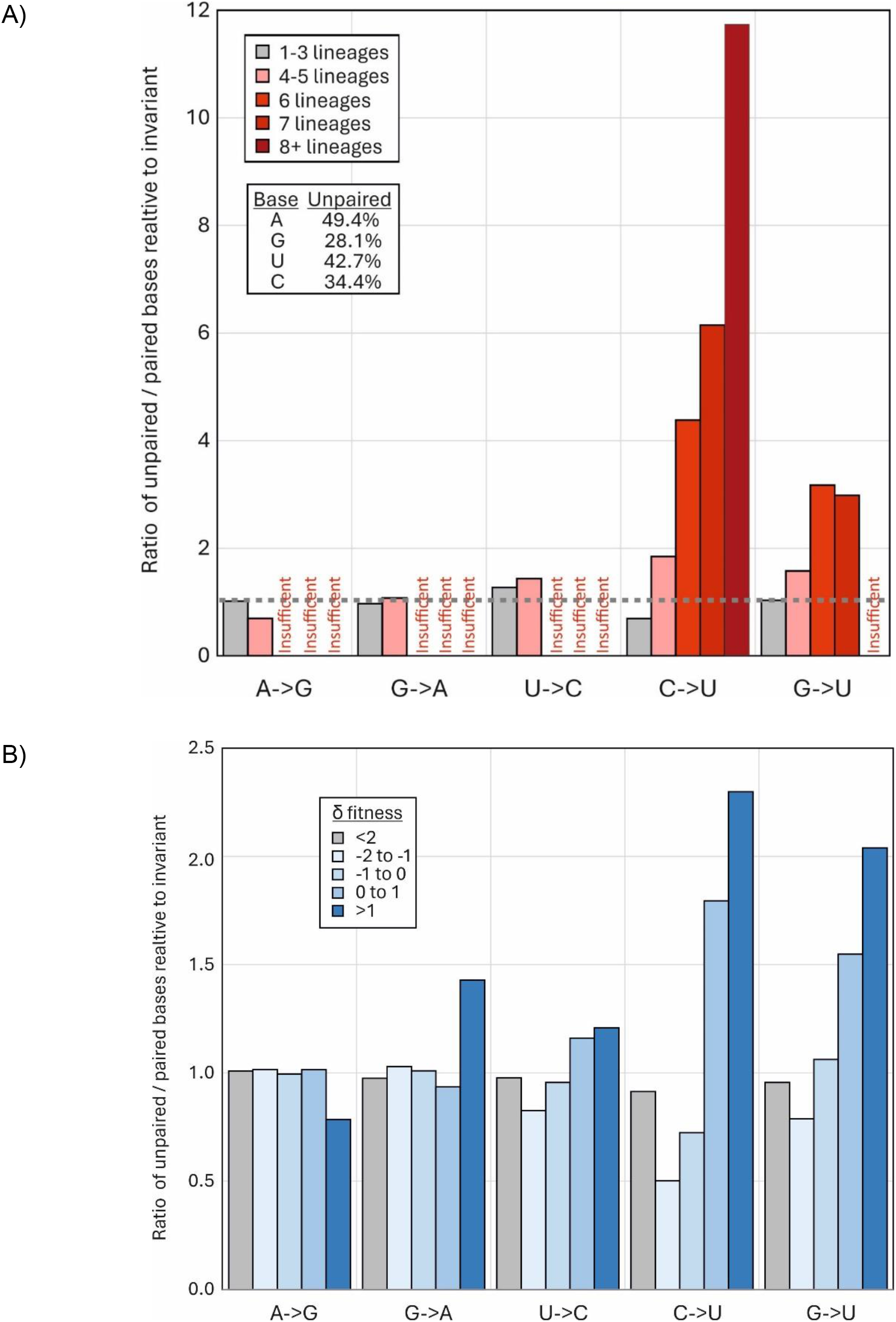
TRANSITION SITE ASSOCIATIONS WITH RNA SECONDARY STRUCTURE PAIRING. A) Comparison of frequencies of unpaired bases at sites with mutations in one or more lineage (grey bars) and in 4, 6, 8 or more lineages (pink, red and dark red bars) with those of invariant sites (proportions of unpaired bases shown in inset table). There were insufficient numbers of lineages with 6 or more A->G, G->A or U->C mutations (<10) to plot. (B) Frequency of unpaired bases at sites classified by the fitness metric, δ*f* (log ratio of observed substitutions / expected number) (15)

Unexpectedly, G->U transversion showed a similar preference for unpaired bases. Frequencies of unpaired sites were similarly higher at more polymorphic sites of C->U transitions in the published analysis of 7 million SARS-CoV-2 sequences (Fig. 8B).

However, as for 5’U context preferences, the majority of C->U over-represented sites (polymorphic in 4+ lineages) did not show a preference for unpaired bases (Table S2B; Suppl. Data).

## DISCUSSION

### Trajectory of SARS-CoV-2 evolution

The publication of an unprecedented number of accurate SARS-CoV-2 complete genome sequences four years on from the start of the COVID-19 pandemic provides an opportunity to re-investigate the trajectory of SARS-CoV-2 evolution. In particular the sequence data enables further exploration of the reported mutational biases driving the high rate of C->U transitions in consensus sequences of SARS-CoV-2 isolates (5–11). The longer timescale provides a convincing demonstration of the sustained loss of C bases and accumulation of Us resulting from this transition bias, with net rates of -0.25% and +0.25% / decade respectively (Figs. 3, 5). These observations provide an insight into the origin of the much greater imbalance of complementary bases (G > C, U > A) in other human seasonal coronaviruses (23, 24) and tentative minimum timescales for their proposed original zoonotic spread into humans. If these originate from C->U hypermutation, the observed base imbalances in the betacoronaviruses HCoV-OC43 and HCoV-HKU1 would predict origins of a minimum of 400 or more years ago if the trajectory of C loss in SARS-CoV-2 was indeed sustained for longer periods. While the former date is much earlier than the proposed attribution of HCoV-OC-43 to the Russian “flu” global outbreak in 1892-3 (25, 26), human infection was proposed to have been acquired via cows as an intermediate host, who may have exerted their own pressure on C->U mutations. The extreme transition asymmetry in SARS-CoV-2 after spread into deer (Fig. 5) indeed suggests that ungulates may harbour even more potent mutational drivers than found in human (and carnivore) cells.

Although there is a lack of large datasets of nucleotide sequences from bat-derived sarbecoviruses, and existing strains are relatively more divergent from each other than between SARS-CoV-2 isolates, there was no evidence for a C->U hypermutation based on comparisons with reconstructed ancestral sequences and a 5% directionality threshold (Fig. 5B), nor were there the marked base frequency asymmetries observed in human seasonal coronaviruses (Fig. 5A). These observations suggest that replication in bats is not associated with the same mutational pressure as found in human or other mammalian infections. While potential mechanisms remain controversial, the host comparisons in the current study provide strong evidence that the hypermutation phenomenon is host-derived rather than being an intrinsic property of sarbecovirus and of wider coronavirus replication strategies, or misincorporation frequencies by the coronavirus RdRP. The existence of C->U/U->C transition asymmetries in the mutational spectra of a wide range of other mammalian +strand RNA viruses (18) similarly argues that the phenomenon represents a more general host-driven phenomenon rather than being connected to the specifics of the SARS-CoV-2 replication complex.

### Mutational biases in SARS-CoV-2 genome sequences

Numerous studies document a range of mutational biases in the evolution of SARS-CoV-2. These include ADAR1-mediated editing of G bases to inosine in viral dsRNA template that resolve to G->A substitutions (and theoretically equal frequencies of T->C mutations derived from editing of the complementary strand). Such mutations were originally described within SARS-CoV-2 populations (6, 8, 9), but are less prominent in very large datasets of consensus sequences from different SARS-CoV-2 isolates (*eg.* (15, 27) and in the original studies (5–11). Instead, all datasets record the dominance of the C->U transition and its asymmetry relative to other transitions. There have been some countervailing views on the existence of this mutational bias (28), but typically these are based on observations of within-population polymorphisms where other mutational effects, such as sequencing errors, may contribute significantly to the observed diversity.

The transition asymmetry in SARS-CoV-2 sequences and the proposed wider occurrence of the phenomenon in some but not all vertebrate RNA viruses (18) is evidently a real phenomenon that requires a mechanistic explanation.

### Drivers of C->U hypermutation

The most frequently advanced hypothesis, now with some experimental evidence, is that observed C->U transition bias arises from direct editing of viral RNA sequences by one or more isoforms of the APOBEC family of nucleic acid deaminases (5–11). APOBECs A3D, A3F, A3G and A3H are potent inhibitors of retroviruses, and co-transcriptionally edit transcripts of DNA during reverse transcription of proviral sequence pre-integration (reviewed in (12)). Errors so introduced create severe and permanent replication defects in the progeny virus. APOBEC activity creates an observable depletion of UpC or UpU dinucleotides in genome sequences of a range of DNA viruses, retroelements and some RNA viruses (29, 30). The observed depletion of UpC in seasonal coronaviruses might be similarly interpreted as evidence for the activity of other APOBEC isoforms, notably human A3A, on RNA sequences. There is also, for example, evidence for extensive A3A-mediated editing of human mRNA sequences as part of a cellular stress response, with targeting of unpaired bases in RNA secondary structures (31, 32), similar to the originally described editing of the ApoB mRNA 3’UTR mRNA by A1 (33, 34). The RNA genome of rubella virus shows a marked C->U/U->C transition asymmetry and genome-wide depletion of UpC (35). The replication of the seasonal coronavirus, HCoV-NL63, was impaired in cell expressing A3C, A3F, and A3H isoforms of APOBEC although in this case it was not demonstrated to result from introduced mutations in the genome (36).

Several studies have documented extensive secondary structure formation in the SARS-CoV-2 positive sense genomic RNA, manifested through the formation of sequential extensively internally base-pair stem-loops throughout the non-coding and coding regions of the genome (20–22, 37). On the face of it, there is therefore no reason to suppose that SARS-CoV-2 genomic RNA or mRNAs might not also be targeted by APOBEC in an RNA structure-dependent manner comparable to what has already been documented for cellular mRNAs and other RNA viruses with single-stranded RNA genomes. Recently, it was demonstrated that over-expression of A3A (but not other isoforms of human APOBEC) induced mutations in the genome of SARS-CoV-2 during *in vitro* passage in 293T cells at several genomic sites (19). These occurred in the favoured 5’U context and occurred in unpaired bases in the terminal loops of RNA secondary structures. Induced mutations did not impair overall replication fitness of SARS-CoV-2, consistent with a reported lack of correlation between extent of C->U editing and SARS-CoV-2 titre on *in vitro* passaging (38). Indeed, another study showed that a functional A3A appeared to enhance replication, perhaps by providing greater opportunities for fixation of phenotypically advantageous mutations, such as C241U in the 5’UTR (39). However, high multiplicity passage would rapidly drive out even marginally deleterious C->U mutations through natural selection and might hide the actual impact of APOBEC-mediated editing on replication fitness.

While the *in vitro* studies provide important functional characterisation of A3A-mediated RNA editing, it is not clear whether A3A drives the bulk of C->U transitions *in vivo*. None of the published datasets analysing consensus sequences of SARS-CoV-2 variants demonstrate a pronounced preference for a 5’U context at sites of C->U transitions that could not be simply accounted for by high genome content of U (and A) bases in the SARS-CoV-2 genome. This includes the set of C->U substitutions fixed during lineage diversification (Fig. 1), and unfixed mutations within lineages (Fig. 5) – for the latter, a modelling approach in which randomisation was applied under same constraints that might operate *in vivo* (coding, and preservation of dinucleotide frequency biases) resulted in 5’ and 3’ base frequencies identical to those surrounding C->U edited sites (Fig. 7A). In this bioinformatic analysis (Fig. 7B) and in the functional studies, it appears that only the most frequently mutated sites *in vivo* or *in vitro* conform to the expected targeting preferences of A3A for a 5’U and positioning inside unpaired regions of stem-loops. However, these represent an extremely small fraction of the total number of *in vitro* edited sites of C->U transitions (19) and those showing elevated C->U transition frequencies in SARS-Cov-2 isolates (Fig. 7B). For example, there was little or no 5’base preference among the unfixed C->U substitutions within 8 or fewer lineages in the current study data (Fig. 7B), corresponding to around 80% of the excess C->U substitutions over other transitions. Similarly, the specific targeting of unpaired bases by A3A recorded in *in vitro* studies was not was not reproduced in site heterogeneity of SARS-CoV-2 isolates (Fig. 8A, 8B). While a higher proportion of highly polymorphic C->U sites were unpaired, a substantial proportion of the excess C->U substitutions also occurred in base-paired regions.

While it has been argued that the lack of context at many C->U mutated sites rules out an editing role for A3A or other APOBEC (28), previous functional investigations of A3A induced changes in SARS-CoV-2 and human mRNA similarly also found variability in editing contexts. These suggest that, unlike APOBEC-mediated editing of DNA sequences, specific contexts may be preferred but not absolutely required. A lack of stringency was suggested by the observation that while C bases in human mRNA were preferentially edited within unpaired regions of stem-loops, around a third of mutated sites occurred in other RNA structure contexts (32). There was similar variability in 5’ contexts and pairing in sites edited in an *in vitro* RNA expression system of a short section of the SARS-CoV-2 genome (39).

### Biological impact of C->U mutations in COVID-19

APOBECs have evolved as a potent and essential defence against many virus infections; the expansion of the A3 locus in primates is thought to have been a host response to the spread of retroviruses early in their diversification (40, 41). Consequently, APOBEC repertoires are highly diverse among different orders of mammals and the existence and activity of precise homologues of A3A (that appears to mediate editing of SARS-CoV-2 in human cells) in other species is poorly understood. Speculatively, the remarkable editing of SARS-CoV-2 in deer (Fig. 4) potentially mediated by one of the much more restricted number of A3 genes in artiodactyls (one with the A3Z1 domain organisation corresponding to human A3A; (42)) perhaps represents an example of a more active restriction mechanism for RNA viruses than found in the human or other primate genome.

RNA viruses with single stranded genomes typically follow the first of Chargaff’s rules that genome frequencies of C should match those of G and A ≈ T/U; these reflect the symmetry of base incorporation biases on copying plus and minus strands of genomic RNAs. However, genome compositions of seasonal coronaviruses violate this principle (Fig. 4; (24)) and it might be speculated that their composition represents a much longer-term endpoint of the loss of C bases and accumulation of U’s apparent in the SARS-CoV-2 genome (Fig. 3).

Contrastingly, bat sarbecovirus show minimal imbalances (Fig. 4) and no evidence for excess C->U substitutions in their diversification (Fig. 5B). These observations suggest either bat APOBECs are incapable of editing sarbecovirus sequences, or more likely, that sarbecoviruses have evolved antagonists to bat APOBECs that prevent genome editing as part of a longer process of virus / host co-adaptation. This might be analogous to the evolution of Vif and other antagonists of APOBECs that target post-entry reverse transcription of retroviral genomic RNA (43, 44). An inability of SARS-CoV-2 to antagonise human A3A is indeed a plausible outcome of the broad genetic and functional diversity of the APOBEC locus in different mammalian orders. As another possible analogy, monkeypox virus (MPXV) infection in humans was associated with the appearance of a clade bearing multiple (40+) and in this case symmetric C->T/G->A mutations attributed to genome editing of the dsDNA genome associated with a failure to antagonise DNA-targeting A3s such as A3G (45). Similarly, an inability of Vif in simian and feline lentiviruses to antagonise APOBECs in heterologous species represented a key factor limiting their host range (46, 47). These examples support a more general hypothesis that the failure of viruses in the “wrong” host to antagonise APOBEC editing pathway may be key determinants in their host range (48).

This background provides some context for understanding the burning question of whether the observed C->U mutations introduced into the SARS-CoV-2 genome during the pandemic have measurably affected its replication and ability to transmit. Related to that, are seasonal coronaviruses irreversibly damaged by their current skewed genome composition (Fig. 4)?

Kim *et al*. proposed that C->U editing may be beneficial by providing a source of novel mutations that might enhance the ability of SARS-CoV-2 to adapt to human transmission and escape from host immune responses (39). Although not functionally mapped, C->U substitutions have been fixed in 68 different sites within the 10 lineages of SARS-CoV-2 analysed in the current study (Fig. 1), consistent with the potential contribution of some of these to replication fitness and increased transmissibility of SARS-CoV-2 variants emerging during the pandemic. On the other hand, the vast majority of C->U substitutions are likely unobserved because of their adverse effect on replication phenotypes; alternatively, they may impart a marginal fitness loss that contributes to observed cycles of mutation and reversion at favoured sites for C->U mutations (9, 18, 39, 49). Unfortunately, while extensive analysis of C->U substitution dynamics using human-derived SARS-CoV-2 strains in this and previous studies provides some insights, there is no real “negative control” to address the question of whether SARS-CoV-2 replication fitness, infectivity and evolutionary rates in natural chains of human-to-human transmission would be enhanced in the absence of C->U hypermutation. The human A3A gene is insufficiently polymorphic in humans to enable such comparisons to be made observationally, and mouse or hamster models with their much more limited host A3 repertoires and functionally quite different APOBEC restriction systems are unlikely to provide an accurate model of human antiviral responses. However, determining the importance of APOBEC-mediated restriction of SARS-CoV-2 and indeed other RNA viruses showing C->U hypermutation is an important future path of investigation that will illuminate the role of this of this pathway in RNA virus defence and in determining host range and zoonotic potential.

